# MAT2A inhibition in AML unveils therapeutic potential of combining DNA demethylating agents with UPR targeting

**DOI:** 10.1101/2023.06.05.543499

**Authors:** Keti Zeka, Alice V. Taylor, Ralph Samarista, Denise Ragusa, Chun-Wai Suen, Oliwia W. Cyran, Ana Filipa Domingues, Eshwar Meduri, Brian JP Huntly, Dorian Forte, Antonio Curti, Cristina Pina

**Author notes:** These authors contributed equally to this work (Keti Zeka, Alice V. Taylor, Ralph Samarista).

## Abstract

Acute Myeloid Leukaemia (AML) is a heterogeneous disease of dismal prognosis, with vulnerabilities in epigenetic and metabolic regulation. DNA demethylating agents, e.g. azacytidine (AZA), are used as first-line therapy in AML patients unable to tolerate intensive chemotherapy regimens, often in combination with BCL-2 inhibitor venetoclax. However, the impact on survival is limited, indicating the need for alternative therapeutic strategies. Methyl-group usage for epigenetic modifications depends on methionine availability and MAT2A-driven conversion to S-adenosyl-methionine. Methyl-group production is a vulnerability in multiple tumours, including AML, and has been variably linked to impairment of different histone methyl-modifications. In contrast, we herein align MAT2A effects in AML with DNA methylation and proteostasis. We show that MAT2A inhibition can be mimicked by combining AZA with unfolded protein response (UPR) activation through targeting of valosin-containing protein (VCP)/P97. Combined AZA and P97 inhibition exceeded AZA-driven restriction of human AML cell expansion, and specifically impaired colony-formation and maintenance of CD34+ patient blasts, suggesting targeting of AML stem/progenitor-like cells. Overall, our data support combined targeting of DNA methylation and the UPR as a promising therapeutic strategy in AML.

---

Acute Myeloid Leukaemia (AML) is a heterogeneous disease of dismal prognosis, with identified vulnerabilities in epigenetic regulation that can be targeted therapeutically. DNA demethylating agents are a therapeutic mainstay in myelodysplastic syndromes and have recently gained traction as a first-line therapy, often in combination with the BCL-2 inhibitor venetoclax [1], in the large group of elderly patients with AML unable to tolerate intensive chemotherapy regimens [1]. However, the impact on survival remains limited, with early emergence of resistance and disease relapse, indicating the need for alternative therapeutic agents or further combinatorial strategies. In addition to DNA methylation, other epigenetic modifications, on histones and the catalytic enzymes that deposit these modifications, have been extensively targeted in AML, with modest and subtype-restricted results seen in clinical trials [1]. Attention has also been given to metabolic dependencies, particularly in relation to the common *FLT3* mutations. Methionine restriction, which results in a reduction of methyl group availability due to reduced production of S-adenosyl-methionine (SAM) from methionine, has consistently been associated with reduced proliferation and survival of cancer cells, both in solid tumours [2,3] and more recently in AML [4]. From an epigenetic point of view, SAM depletion differentially affects H3K4me3 [5], H3K27me3 [6] or H3K36me3 [4] depending on tumour type, with the latter more specifically targeted in AML [4].

MAT2A is the enzyme that catalyses SAM synthesis from methionine in most mammalian cell types, including blood cells, and we previously identified it as a genetic vulnerability in a CRISPR screen of AML cell lines (7-summarised in Supplementary Fig. 1A). We verified MAT2A dependence chemically, through treatment with FIDAS-5 (Supplementary Fig. 1B), a potent and orally active MAT2A inhibitor, and by gene expression knockdown (Supplementary Fig. 1C-D). Importantly, we observed that the effects of MAT2A depletion extended to *in vivo* engraftment of AML cells (Fig. 1A-B), with reduced graft contribution of MAT2A knockdown cells (Fig. 1B) and prolonged survival of MAT2AshRNA-engrafted animals (Fig. 1A). Furthermore, depletion of MAT2A impacted colony-formation from AML primary patient samples (Fig. 1C) across multiple mutational backgrounds (Supplementary Table 1), an effect that was selective for AML cells, and not observed in healthy cord blood progenitors (Supplementary Fig. 1E). These observations match a recent study highlighting MAT2A as a vulnerability in patient-derived xenograft (PDX) cell lines carrying *KMT2A/MLL*-rearrangements [8]. RNA-sequencing (RNA-seq) analysis reveals metabolic reprogramming of AML cells with MAT2A inhibition, affecting oxidative phosphorylation, production of reactive oxygen species, adipogenesis and fatty acid metabolism, as well as DNA repair signatures (Fig. 1D). MAT2A and methionine metabolism have been previously characterised as specific dependencies for deposition of Histone 3 Lysine 4 tri-methylation (H3K4me3) in solid tumors [9] and for H3K36me3 in AML [4]. However, our analysis of a small number of promoters and bodies of genes differentially down-regulated upon FIDAS-5 treatment suggest a heterogeneous pattern of histone methylation changes (Supplementary Fig. 1F-G). In addition, inspection of DNA demethylating agent azacytidine (AZA)-associated programs in treated MOLM-13 cells revealed an enrichment of AZA targets amongst MAT2A signature genes (Fig. 1E). Consistently, we observed a global reduction in methylated cytosine residues upon MAT2A loss (Fig. 1F-G), linking the epigenetic effects of MAT2A and methionine to DNA methylation. Furthermore, RNA-sequencing (RNA-seq) analysis of MAT2A-inhibited cells captured up-regulation of elements of the pro-apoptotic arm of the unfolded protein response (Fig. 1D, H), which we followed up on as a novel candidate therapeutic approach currently unexplored in AML.

**Figure 1.**
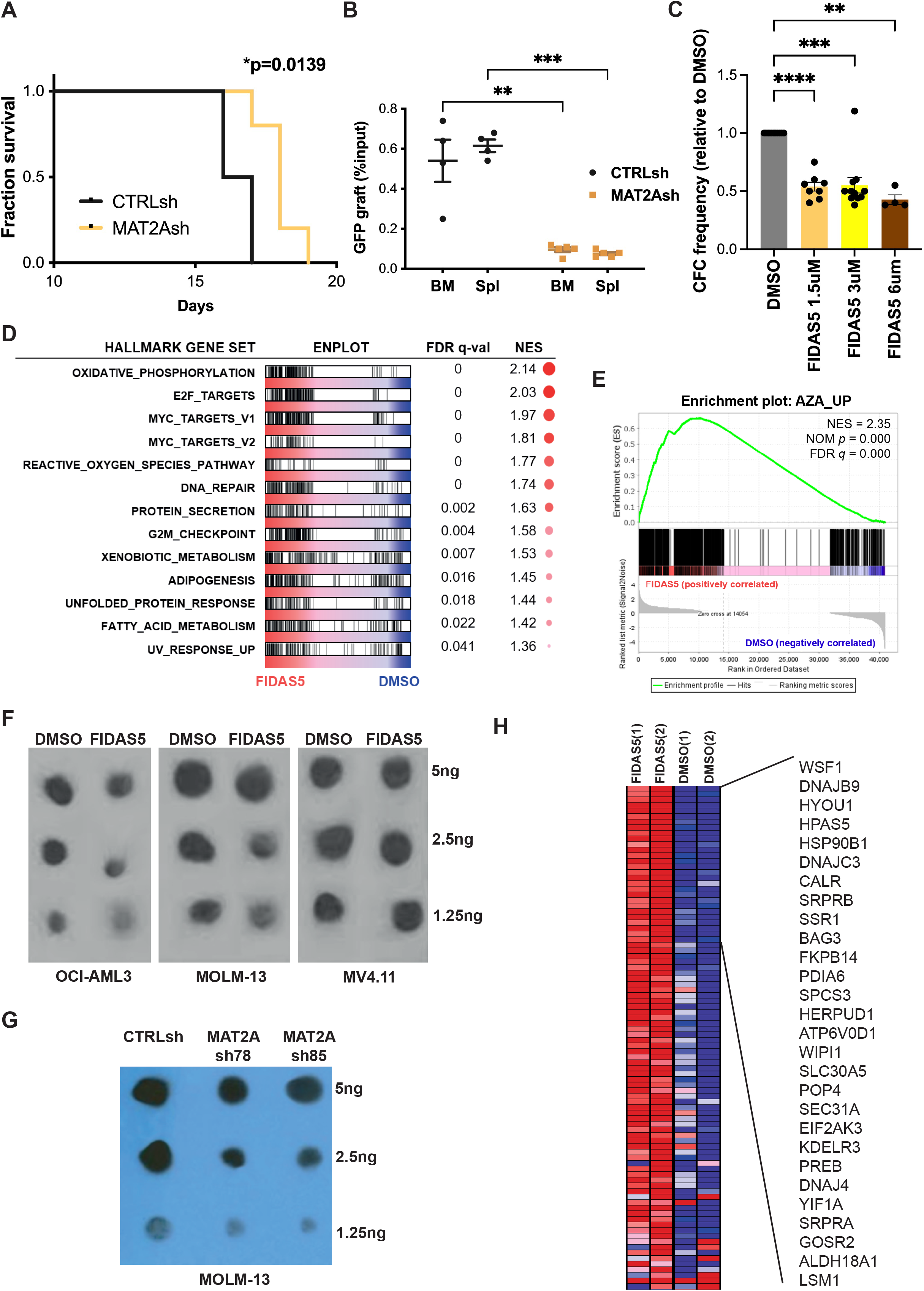
MAT2A loss inhibits AML growth through effects on DNA methylation and proteostasis. **A**. Survival curve of NSG mice transplanted with lentiviral-transduced MOLM-13 cells expressing pooled shRNA against MAT2A; n=5 MAT2Ash recipients; n=4 control, CTRLsh recipients; log-rank test, *p<0.05. **B**. Relative GFP engraftment of CTRL and MAT2Ash recipients at the point of culling. Analysis of engraftment in bone marrow, BM and spleen, plotted relative to input %GFP. Two-tailed t-test, **p<0.01, ***p<0.001. Animals as in A. **C**. Colony-forming cell activity in n=11 AML patient samples treated with 1.5 (n=8), 3 (n=11) or 6μM FIDAS-5 or DMSO (0.1% final). Data relative to DMSO; paired t-test **p<0.01, ***p<0.001, ****p<0.0001. **D**. Hallmark gene set enrichment analysis (GSEA) of RNA-seq data obtained from FIDAS-5 (3μM) vs DMSO-treated MOLM-13 cells after 48h treatment. Enrichments at FDR<0.05 are represented. **E**. GSEA of AZA up-regulated signatures in the FIDAS-5 treatment RNA-seq data described in D. AZA signatures were obtained from MOLM-13 cells treated with 7μM AZA (Cayman Chemicals) for 48h. **F-G**. DNA methylation dot blots of OCI-AML, MOLM13 and MV4.11 cells treated for 48h with FIDAS-5 3μM vs DMSO (0.1% final) (**F**) or lentiviral-transduced MOLM-13 cells (**G**). **H**. Leading-edge gene list from enriched GSEA Hallmark Gene Set ‘Unfolded Protein Response’ in D. The top 30 genes are specified. Red indicates up-regulation of gene expression.

Proteostasis is a recently recognised vulnerability of cancer cells [10] and plays a critical role in haematopoietic stem cell maintenance. Specific activation of the apoptotic arm of the UPR through p97 inhibition has been explored as a therapeutic strategy in B Acute Lymphoblastic Leukaemia (B-ALL) [11] and multiple myeloma [12], as well as in solid tumours [13]. However, its significance in AML is unclear [14]. We initially verified the anti-proliferative effects of UPR activation in 2 different AML cell lines (Supplementary Fig. 2A-B), using 2 different p97 inhibitors (CB-5083, and NMS-873) [14]. We next verified activation of UPR genes in the same cell lines (Supplementary Fig. 2C), and determined the IC50 concentrations (viability) for downstream experiments (Supplementary Fig. 2D). Given the combined effect of MAT2A inhibition on DNA demethylation and UPR activation, we hypothesised that combinatorial targeting of both pathways might have enhanced anti-AML effects, and together capture the observed consequences of MAT2A loss. We started by treating AML cell lines MOLM-13 (Fig. 2A-B) and OCI-AML3 (Supplementary Fig, 2E-F) with the p97 inhibitor CB5803 (CB) and/or AZA at IC50 (full) or half-IC50 (half) concentrations. We observed a significant reduction in cell proliferation for all single and combinatorial treatments relative to vehicle (DMSO) (Fig.2A, Supplementary Fig. 2E). Combinations of full-dose CB with full- or half-dose AZA resulted in significant reductions in cell expansion compared to AZA or CB alone (Fig. 2A, Supplementary Fig. 2E), confirming the potential of combining both drugs. Similar results were obtained when inspecting cell viability in the treated cultures (Fig. 2B, Supplementary Fig. 2F), particularly at the time-points in which combinatorial treatment conferred an anti-proliferative advantage (respectively, 96h for MOLM-13 (Fig. 2B), and 144h for OCI-AML3 (Supplementary Fig. 2F)). We next analysed the effects of CB and AZA treatments on the frequency of colony-forming cells within AML cell cultures, as a surrogate for AML propagation. Again, we observed a significant reduction in colony-formation, suggesting an impact in AML progenitor cells by combinations of full-dose CB with full or half AZA doses (Fig. 2C), in comparison with both control and AZA alone. To determine the translational value of our observations, we tested the combination of CB and AZA in AML patient samples (Supplementary Table 1), investigating both colony formation (Fig. 2D) and the effect on stem/progenitor phenotypic marker CD34 by flow cytometry (Fig. 2E). Repeatedly, combinatorial treatment decreased colony-formation from AML patient blasts to a greater extent than AZA (Fig. 2D), which did not reach significance in isolation. In these patients, CB significantly reduced colony-formation (Fig. 2D), an observation corroborated by the reduction in CD34+ leukemic progenitors in at least one of the patients (Fig. 2E - left panel). While inter-patient variability reflects the heterogeneity of AML, the data nevertheless support the added value of combining demethylating agents with UPR activation in at least a subset of patients currently treated with AZA alone. Additionally, the results support the notion that MAT2A inhibition can be mimicked, at least in part, by targeting these 2 pathways (Fig. 2D). We validated the similarities between combined AZA+CB treatment and FIDAS-5 through RNA-seq gene expression analysis of MOLM-13 cells exposed to individual treatments or to the AZA+CB combination (Fig. 2F-G). We observed considerable overlap between CB and AZA treatments with FIDAS-5, but also amongst AZA and CB themselves, suggesting that they may converge mechanistically (Fig. 2G). AZA and CB signatures nevertheless have unique enrichments, which potentially target non-overlapping leukemic pathways, potentially making a combinatorial approach more robust against drug resistance mechanisms. Combined AZA+CB signatures were more extensively enriched than individual treatment against FIDAS-5-upregulated transcriptional programs (Fig. 2F), confirming the similarity between dual DNA methylation and UPR targeting and MAT2A inhibition.

**Figure 2.**
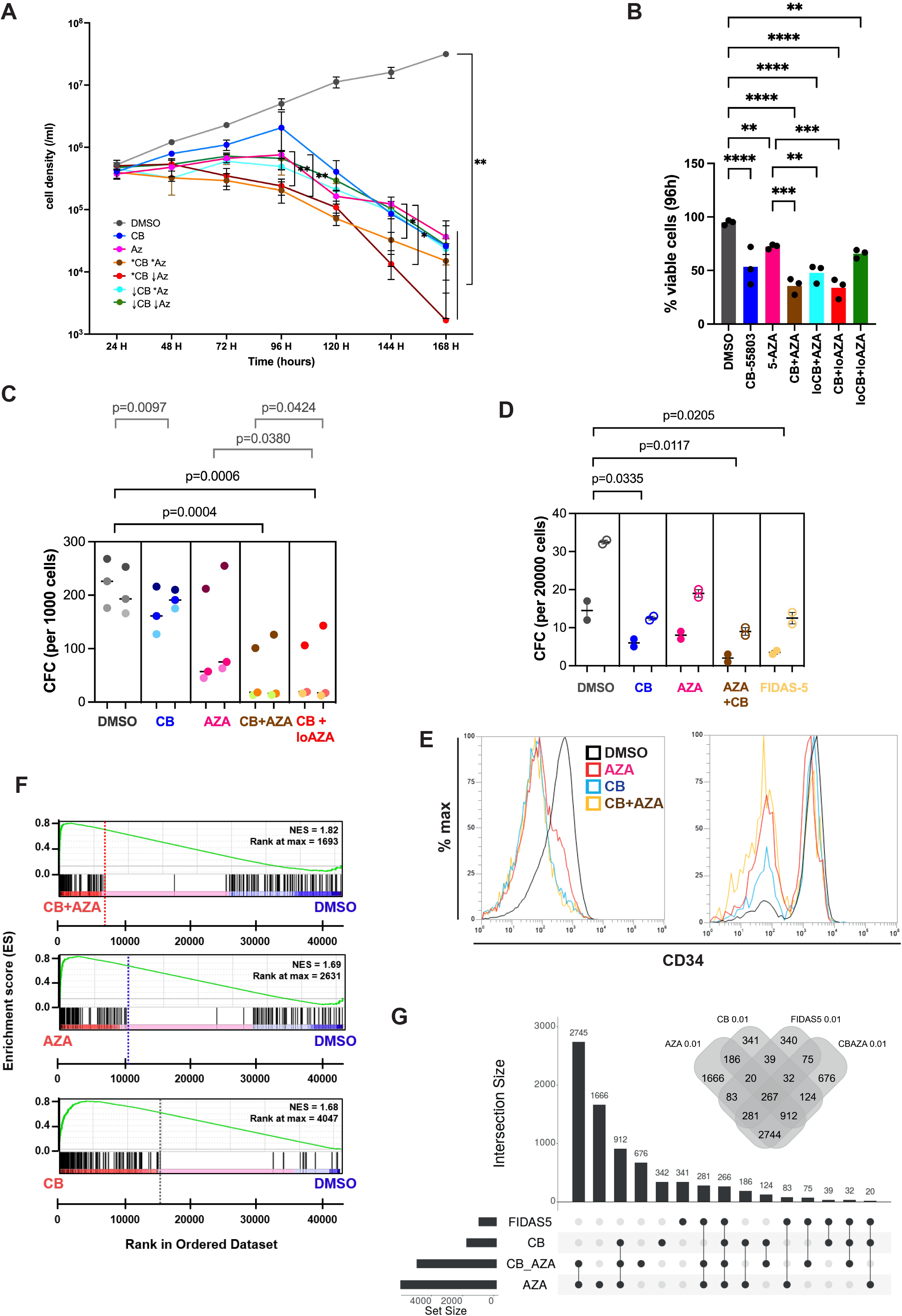
Combined targeting of DNA methylation and the unfolded protein response have therapeutic potential in AML. **A**. Growth curve of MOLM-13 cells treated with CB5083 (CB), AZA or a combination of both compounds in comparison with DMSO (0.1% final volume). CB and AZA were used at IC50 (*) or half-IC50 (down arrow) dose (Supplementary Figure 2D, AZA=7μM). Mean±SD of 3 replicates; mixed-effect analysis with significant effect of treatment (p=0.004); Tukey’s multiple comparisons test, *p<0.05, **p<0.01. **B**. MOLM-13 viability at the 96h-time point of the culture in A. One-way ANOVA (p<0.0001) with Fisher’s LSD multiple comparisons, **p<0.01, ***p<0.001, ****p<0.0001. **C**. Colony-forming assays of MOLM-13 cells treated with full CB5083 and/or AZA IC50 doses (or half IC50 AZA, lo, in combination). N=3 independent replicates performed as technical duplicates; paired t-test. **D**. Colony-forming assays of 2 individual patient samples performed as technical duplicates in the presence of CB5083 (CB, 1μM), AZA (16μM) or FIDAS-5 (3μM) in comparison with DMSO (0.1% total volume). Paired t-test with significant p-values shown. **E**. Flow cytometry plots of patient samples after 24-culture in the presence of CB±AZA in comparison with DMSO (0.1% final). **F**. GSEA plots for FIDAS-5 up-regulated signatures against CB, AZA and CB+AZA vs. DMSO treated RNA-seq analysis MOLM-13 cells. Cells were treated for 48h with IC50 doses of the respective compounds in/or 0.1% DMSO. **G**. Differential gene set overlaps between CB and/or AZA and FIDAS-5 treatment of MOLM-13 cells as per RNA-seq data analysis at FDR<0.01.

In conclusion, we identified a molecular synergism between currently used demethylating agent AZA and the p97 inhibitor CB5083. We showed that the combination has robust anti-leukemic effects in both AML cell lines and AML patient samples of diverse mutational composition, suggesting the potential for broad therapeutic application. Importantly, the combination is more effective than AZA at targeting phenotypic and functional leukemia stem/progenitors, with putative value in achieving sustained responses in patients.

## MATERIALS AND METHODS

### Cell lines

MOLM13, OCI-AML3 and MV4.11 lines were obtained from DSMZ or the Sanger Institute and their immunophenotype was validated against the DSMZ database. Cell lines were maintained in R20 medium consisting of RPMI (Merck or Fisher, UK) supplemented with 20% FBS (Fetal Bovine Serum, Merck, UK), 1% Penicillin/Streptomycin/Amphotericin (P/S/A, Fisher, UK) and 2mM L-Gln (Fisher, UK). HEK 293T cells were obtained from ATCC; cells were grown in DMEM (Merck or Fisher, UK) supplemented with 10% FBS, and P/S/A and L-Gln (D10). Cultures were kept at 37°C and 5% CO_2_. Cell lines were confirmed as Mycoplasma negative at regular intervals during the study. For inhibitor treatment, FIDAS-5 (Abcam, UK), Azacytidine (AZA, Merck, UK), CB-5083 (Cayman Chemical, UK) and NMS-873 (Cayman Chemical) were reconstituted in DMSO (Fisher, UK) as 1000X stock solution and added to the cultures at the indicated concentrations in 0.1-0.2% final volume of DMSO. Cells were counted daily with live/dead cell discrimination in the presence of Trypan Blue (Fisher, UK).

### Human cord blood and AML patient samples

Cord blood samples and AML patient samples were obtained with informed consent under local ethical approval: REC 07MRE05-44 (Cambridge) and EC code number 94/2016/O/Tess (Bologna). Mononuclear cells (MNC) were isolated by Ficoll-Paque density gradient (StemCell Technologies, UK) centrifugation. CB MNC were enriched for CD34+ cells using the RosetteSepTM Human CB CD34 Pre-Enrichment Cocktail (Stem Cell Technologies, UK) as per manufacturer’s instructions. For some experiments, AML patient samples were cultured for 24h in R20 supplemented with human recombinant IL-3, TPO and G-CSF, all at 20ng/ml (Peprotech, UK).

### Colony-forming cell (CFC) assays

Human AML and CB cells were seeded for 10-12 days in methylcellulose-based semi-solid medium (StemMACS HSC-CFU Media complete with EPO, Miltenyi Biotec, Germany) for assessment of the frequency of colony formation. Cells were resuspended in 10% final volume of R20 prior to addition to the semi-solid medium. Assays were plated in duplicate onto 35mm-dishes or in the wells of a 6-well plate. In the case of CFC assays performed in the presence of an inhibitor, this was added to the methylcellulose-based media and dispersed by vortexing, prior to addition of the cells.

### Lentiviral packaging and transduction

Viral constructs containing control shRNA (CTRLsh, non-eukaryotic gene targeting – target sequence: *CAACAAGATGAAGAGCACCAA*) or shRNA targeting MAT2A (MAT2Ash78 – target sequence: *GTTCAGGTCTCTTATGCTATT*; MAT2Ash85 – target sequence *AGCAGTTGTGCCTGCGAAATA*) in a pLL3.7 backbone (Addgene#11795) were packaged in HEK 293T cells as previously described [15] using Turbofect (Fisher, UK) or trans-IT (Mirus. USA) as lipofection reagents. Cell lines were transduced overnight with 1-2 T75 packaging flask-equivalents (FE)/10^6^ cells and washed the following day, also as described (15). Cells were analysed or sorted for GFP^+^ cells 2-4 days after transduction.

### Transplantation of immunodeficient animals

NOD.Cg-Prkdc scid Il2rg tm1Wjl /SzJ (NSG) mice were purchased from Charles River Laboratories (UK) and housed in a pathogen-free animal facility with unrestricted access to food and water. Experiments were conducted in accordance with UK Home Office regulations, under a UK Home Office project license. Half-million MOLM-13 cells transduced with CTRLsh or pooled MAT2Ash lentiviral particles were injected into sub-lethally-irradiated (2Gy) animals 24h after transduction. Five-percent cells were kept in culture for determination of %input GFP+ 48h later. Animals were monitored daily and were culled upon signs of disease as per the approved experimental plan. Bone marrow and spleen were collected, processed to single-cell suspension, and analysed by flow cytometry for GFP^+^ cells.

### Flow cytometry

Cells were analysed by flow cytometry for GFP expression or cell surface staining using Gallios (Beckman Coulter, UK), Attune (Thermo, UK) or Novocyte (ACEA, UK) instruments. Phenotyping of AML patient samples used a 1:50 dilution of the PE-Cy7-conjugated anti-human CD34 antibody (clone 581, Biolegend, UK). Samples were stained, washed, and resuspended in medium or in 1x MoJo buffer (Biolegend, UK).

### DNA methylation dot blot

Genomic DNA from AML cell lines was obtained by phenol-chloroform-isoamyl alcohol extraction and quantified on a NanoDrop spectrophotometer (Thermo Scientific, UK). DNA was denatured at 99°C for 5 min, cooled quickly on ice and spun down before use. Specified amounts of DNA were blotted on a Hybond N+ membrane (GE Healthcare, UK) in a 1μl volume and left to air-dry. The membrane was blocked in 5% milk in TBS-T for 1 hour at room temperature (rt), with gentle shaking. The membrane was incubated with a 5-mC antibody (GT4111, GeneTek, UK) at 1:500 for 2 hours (rt), followed by 3×10min washes with TBS-T (rt). The membrane was then incubated with anti-mouse IgG HRP-conjugated secondary antibody () at 1:10 000 in 5% milk in TBS-T for 1 hour (rt). The membrane was washed 3×10mins with TBS-T and 1×10mins with TBS before developing the signal by incubating the membrane in Amersham Hyperfilm ECL for 5 minutes (GE Healthcare, UK).

### Western blotting

Western blotting was performed as described [15]. Primary antibodies used were anti-MAT2A (A304-279A, Bethyl Laboratories, UK) and anti-tubulin (sc-5286, Santa Cruz Biotechnologies, USA).

### Chromatin immunoprecipitation (ChIP)

Chromatin was prepared in duplicate from MOLM-13 cells (6 sonication cycles, 30”ON/30”OFF) and immunoprecipitated using anti-H3K4me3 (Ab8580, Abcam, UK) or anti-H3K36me3 (C15310058, Diagenode, France), as described [15]. Eluted DNA was diluted and quantified by SYBR green qPCR using 2μl DNA per triplicate reaction. Peak enrichments relative to input were determined using the 2^-ΔΔCt^ method in reference to *KRT5* locus. Primers used were:

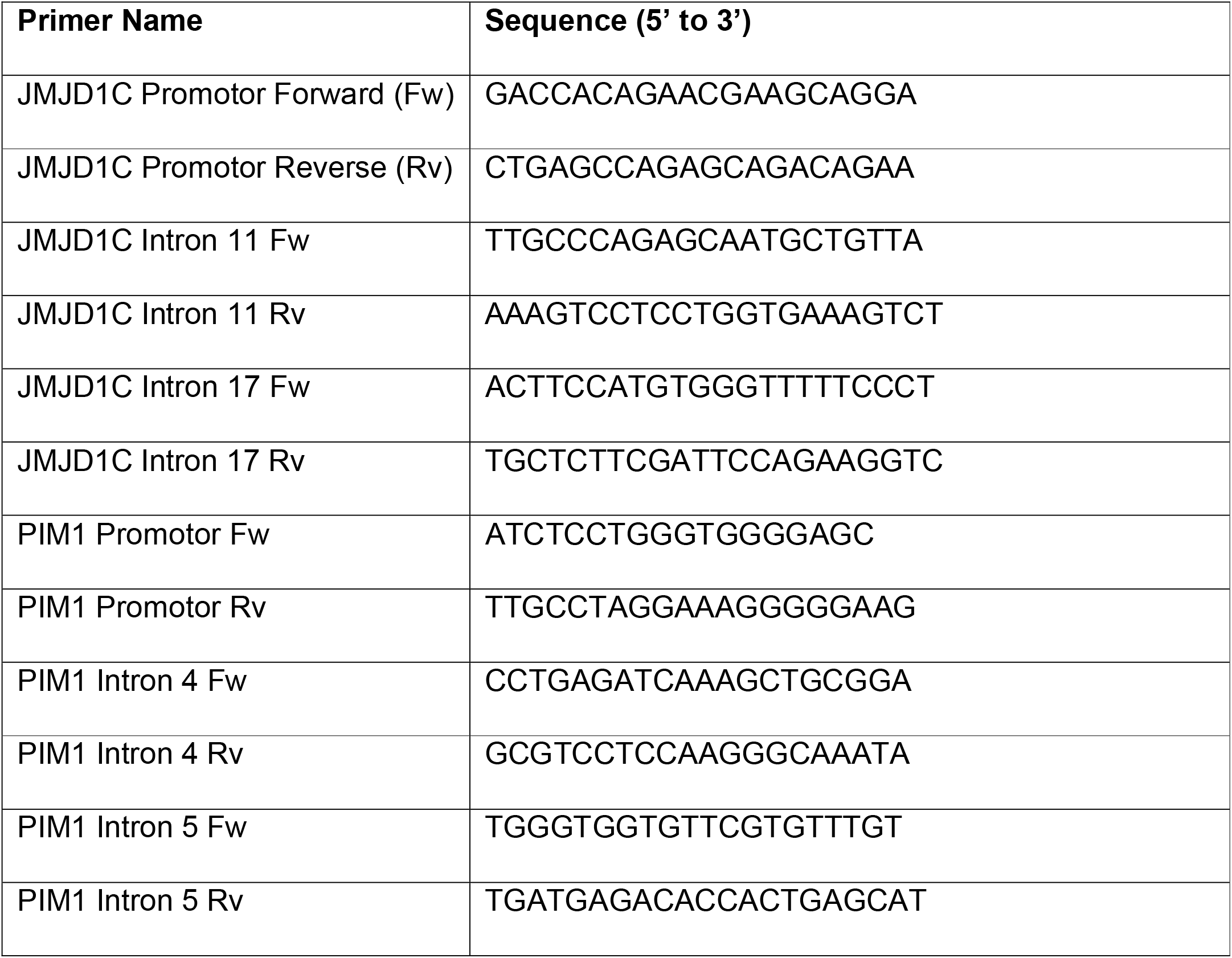
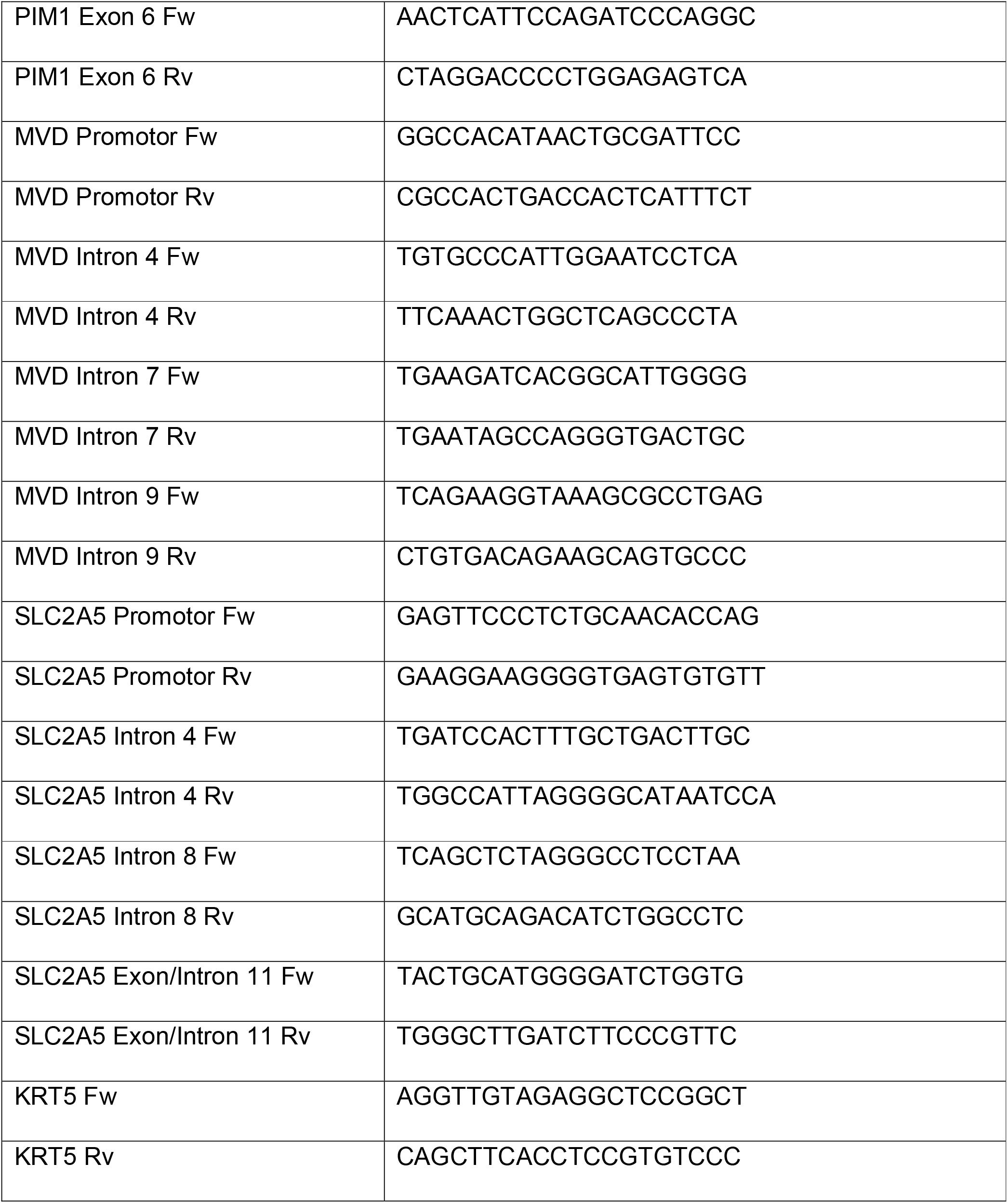

### Quantitative (Q)RT-PCR

RNA extraction, cDNA synthesis and Q-PCR analysis were performed as described [15]. Relative gene expression calculated by the 2^-ΔΔCt^ method using *HPRT1* as reference. Primers used were:

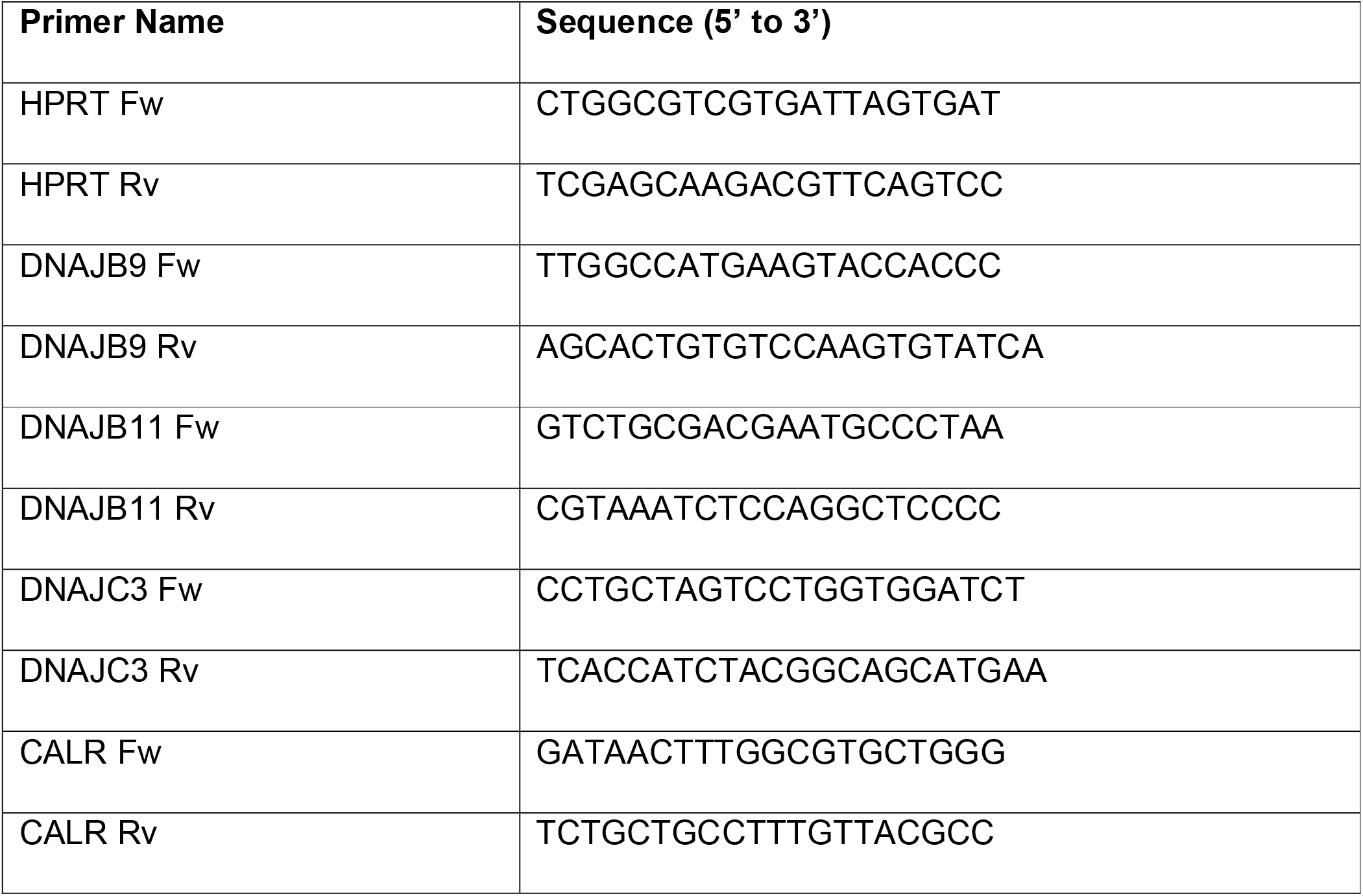

### RNA sequencing

Total RNA from duplicate samples of MOLM-13 cells treated for 48h with FIDAS5 (3μM) or 0.1% DMSO was used for RNA-seq library preparation with ribo-depletion at the Cambridge Stem Cell Institute Genomics Core Facility. Libraries were sequenced on an Illumina HiSeq4000 instrument at the CRUK Cambridge Research Institute Genomics Core Facility using 50bp single-end reads. Total RNA from triplicate samples of MOLM-13 cells treated for 48h with CB5083 (450nM), AZA (7μM), both, or with 0.1% DMSO, were used for polyA RNA-seq library preparation at commercial provider Novogene. Libraries were sequenced on an Illumina NovaSeq PE150 instrument using 150bp paired-end reads. The raw fastq files were mapped using Tophat2 with Bowtie2 and aligned to the reference human genome assembly GRCh38. Differentially expressed genes were obtained using the *Deseq2* R library (v 1.32.0) and filtered by FDR. Hallmark gene sets used for gene set enrichment analysis (GSEA) were obtained from MSigDB and ran on GSEA software v4.2.3 using Signal2Noise metric.

### Statistical analysis

Data were analysed using GraphPad Prism (version 9.5.1.), with the individual tests described in the respective figure legends. IC50 calculations used the Quest Graph™ IC50 Calculator (AAT Bioquest, Inc., 2021) (https://www.aatbio.com/tools/ic50-calculator).

## Data availability

RNA-seq data were deposited in EBI Biostudies under accession E-MTAB-12847 (FIDAS-5) and E-MTAB-12848 (single and combinatorial CB and AZA treatments).

## ACKNOWLEDGMENTS

This work was funded by a Leuka John Goldman Fellowship for Future Science (2017-2019), a Wellcome Trust/University of Cambridge ISSF grant (2019), and a British Society for Haematology Early-Stage Grant (2020-2022) to CP. KZ was funded by a MSCA Post-Doctoral Fellowship (2017-800274). D.F. was supported by Associazione Italiana Ricerca sul Cancro (AIRC-Fellowship 20930 for Abroad). EM was funded by an European Research Council Consolidator Grant (647685) awarded to BJPH. BJPH was also supported by the NIHR Cambridge Biomedical Research Centre (BRC-1215-20014) and Wellcome Trust who funded the Wellcome – MRC Cambridge Stem Cell Institute (203151/Z/16/Z).

## CONTRIBUTIONS

CP conceived and designed the study; KZ, AVT, RS, CS, OWC and AFD collected and assembled data; DF, BJPH and AC contributed critical reagents; KZ, AVT, RS, DR, CS, EM and CP analysed data; KZ, RS, DR and CP interpreted data; KZ and CP wrote the manuscript. All authors approved the final version of the manuscript.

Correspondence: Cristina Pina, College of Health, Medicine and Life Sciences, Division of Biosciences, Brunel University London, Uxbridge UB8 3PH, United Kingdom;

## ETHICS DECLARATIONS

Human cord blood and patient samples were obtained with informed consent under local ethical approval: REC 07MRE05-44 (Cambridge) and Ethical Committee approval code number 94/2016/O/Tess (Bologna).

All animal experiments were conducted in accordance with UK Home Office regulations, under a UK Home Office project licence (PPL80/2454).

## SUPPLEMENTARY MATERIAL

**SUPPLEMENTARY TABLE 1: AML Patient sample information**. Summary of clinical and pathological characteristics of the AML blast samples in this study.

## SUPPLEMENTARY FIGURES

**Supplementary Figure 1. MAT2A is required for maintenance of AML cell lines. A**. Summary of CRISPR drop-out screen results in (8). **B**. Fold expansion of MOLM-13, MV4.11 and OCI-AML3 cells after 3-day culture in the presence of a range of FIDAS-5 doses (or DMSO, 0, 0.1% final volume). Mean±SD of 3 replicates; ANOVA with significant dose effect (p=0.0073); Tukey’s multiple comparisons test, *p<0.05, **p<0.01. **C**. Western blot of MAT2A knockdown in OCI-AML3 (shMAT2A78) and MOLM-13 cells (shMAT2A85). **D**. Growth curves of MOLM-13, OCI-AML3 and MV4.11 cells transduced with lentiviral vectors encoding MAT2A or control (shLLX)-targeting shRNA constructs. **E**. Colony-forming activity of cord blood progenitors treated with 3 different doses of FIDAS-5 or DMSO (0.1% final volume). Mean±SD of 3 individual cord blood samples relative to DMSO; paired-t-test. **(F-G)** ChIP-qPCR analysis of H3K4me3 (**F**) and H3K36me3 (**G**) marks on *JMJD1C, MVD, PIM1* and *SLC25A* loci. Results are shown for 2 independent experiments, with data represented relative to chromatin input and normalised to the unrelated *KRT5* locus. The heatmap panel summarises the average of the two independent experiments for overall trend.

**Supplementary Fig. 2. Targeting of the unfolded protein response affects growth and viability of AML cell lines. A-B**. Growth curves of MOLM-13 (**A**) and OCI-AML3 (**B**) cells treated with serial concentrations of CB5083 and NMS-873 for 72h; 0=DMSO, 0.1% final volume. Mean±SD of 3 replicate experiments; 2-way ANOVA with dose effect p<0.0001 in the 4 treatment / cell line conditions; Dunnett multiple comparison’s test *p<0.05, **p<0.01, ***p<0.001. **C**. Quantitative RT-PCR analysis of *CALR, DNAJB9, DNAJB11* and *DNAJC3* in MOLM13 and OCI-AML3 lines treated with CB5083 or NMS-873 at IC50 values (D) for 48h. Mean±SD of 3 replicate experiments; 2-way ANOVA with treatment effect p<0.0001; Fisher LSD multiple comparisons *p<0.05, *p<0.01, *p<0.001, *p<0.0001. **D**. Calculation of 50% inhibitory concentration (IC50) for effects of CB5083 and NMS-873 on viability of MOLM-13 and OCI-AML3 cell lines. Analysis of 3 independent replicates after 72h of culture. **E**. Growth curve of OCI-AML3 cells treated with CB5083 (CB), AZA or a combination of both compounds in comparison with DMSO (0.1% final volume). CB and AZA were used at IC50 (*) or half-IC50 (down arrow) dose (D, AZA=8μM). Mean±SD of 3 replicates; mixed-effect analysis with significant effect of treatment (p=0.004); Tukey’s multiple comparisons test, *p<0.05, **p<0.01. **F**. OCI-AML3 viability at the 144h-time point of the culture in E. One-way ANOVA (p<0.0001) with Fisher’s LSD multiple comparisons, **p<0.01, ***p<0.001, ****p<0.0001.

