## Supplementary figures and images for "MAT2A inhibition in AML unveils therapeutic potential of combining DNA demethylating agents with UPR targeting"

### Supplementary Figure 1

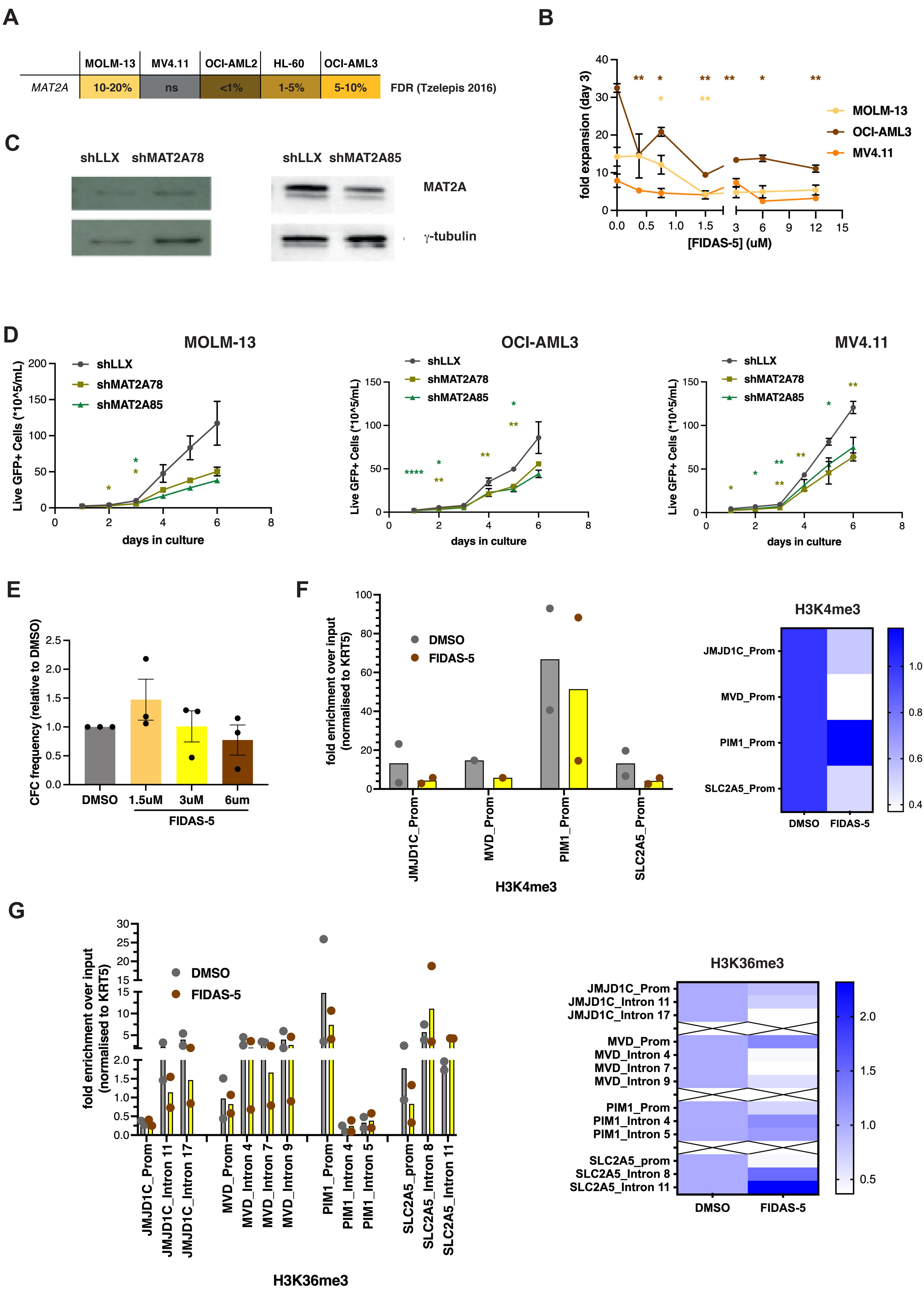

### Supplementary Figure 2

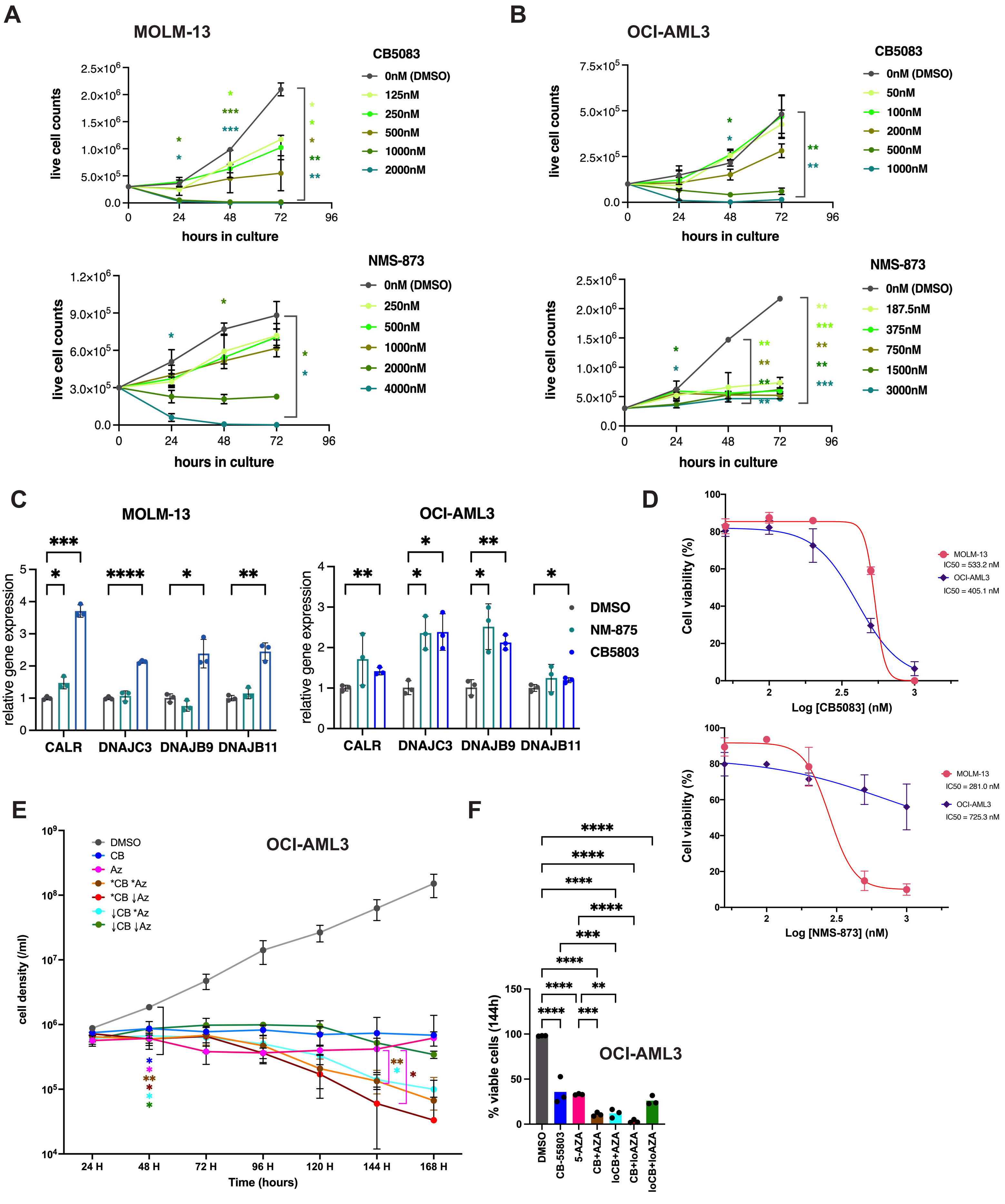
